# Asymmetric dispersal is a critical element of concordance between biophysical dispersal models and spatial genetic structure in Great Barrier Reef corals

**DOI:** 10.1101/453001

**Authors:** C Riginos, K Hock, AM Matias, PJ Mumby, MJH van Oppen, V. Lukoschek

## Abstract

**Aim:** Widespread coral bleaching, crown-of-thorns seastar outbreaks, and tropical storms all threaten foundational coral species of the Great Barrier Reef, with impacts differing over time and space. Yet, dispersal via larval propagules could aid reef recovery by supplying new settlers and enabling the spread of adaptive variation among regions. Documenting and predicting spatial connections arising from planktonic larval dispersal in marine species, however, remains a formidable challenge.

**Location:** The Great Barrier Reef, Australia

**Methods:** Contemporary biophysical larval dispersal models were used to predict longdistance multigenerational connections for two common and foundational coral species (*Acropora tenuis* and *Acropora millepora*). Spatially extensive genetic surveys allowed us to infer signatures of asymmetric dispersal for these species and evaluate concordance against expectations from biophysical models using coalescent genetic simulations, directions of inferred gene flow, and spatial eigenvector modelling.

**Results:** At long distances, biophysical models predicted a preponderance of north to south connections and genetic results matched these expectations: coalescent genetic simulations rejected an alternative scenario of historical isolation; the strongest signals of inferred gene flow were from north to south; and asymmetric eigenvectors derived from north to south connections in the biophysical models were significantly better predictors of spatial genetic patterns than eigenvectors derived from symmetric null spatial models.

**Main conclusions:** Results are consistent with biophysical dispersal models yielding approximate summaries of past multigenerational gene flow conditioned upon directionality of connections. For *A. tenuis* and *A. millepora*, northern and central reefs have been important sources to downstream southern reefs over the recent evolutionary past and should continue to provide southward gene flow. Endemic genetic diversity of southern reefs suggests substantial local recruitment and lack of long distance gene flow from south to north.

## Introduction

Recurrent mass bleaching events on the Great Barrier Reef (GBR) have been increasing in severity and extent (Hughes et al., 2017) against a backdrop of multidecadal coral decline arising from tropical storms, crown-of-thorns seastar (COTS) predation, terrestrial runoff, and fishing pressure (De’ath, Fabricius, Sweatman, & Puotinen, 2012). Locations of severe impacts, however, differ over time. For example, northern reefs have been less affected by water quality problems and COTS outbreaks (De’ath et al., 2012), but yet the highest rates of bleaching in 2016 (the most recent and extensive mass bleaching event) were reported from these reefs (with declines of up to 60 % of coral cover: Hughes et al., 2018). Within the GBR, central reefs have been most affected by terrestrial runoff, episodic COTS outbreaks (Pratchett, Caballes, Rivera-Posada, & Sweatman, 2014), tropical storms, and bleaching (De’ath et al., 2012). Southern reefs also have been subject to COTS outbreaks and tropical storms (De’ath et al., 2012), yet largely escaped bleaching in 2016 (Hughes et al., 2018).

Corals, like most benthic marine animals, have planktonic larvae potentially capable of extensive dispersal. External supplies of settlers can replenish populations; for example, following local extirpation of mature *Acropora* colonies by Cyclone Yasi, recruitment of juvenile *Acropora* was high (Lukoschek, Cross, Torda, Zimmerman, & Willis, 2013). Dispersal connections arising from planktonic larval movements can also enable the spread of adaptive variation. With temperatures and extreme heating events projected to increase in frequency (Wolff, Mumby, Devlin, & Anthony, 2018), resolving the GBR-wide capacity for adaptive gene flow (especially involving loci contributing to heat tolerance) will contribute to emerging debates regarding assisted migration and genetic rescue (Anthony et al., 2017). Of particular interest is to uncover routes of natural connections as well as pathways resistant to gene exchange.

Unfortunately, dynamics of planktonic larval dispersal over space and time remain poorly resolved (Carr et al., 2003; Kritzer & Sale, 2004) especially considering an extensive and geographically complex seascape such as the GBR. Currents and other oceanographic phenomena are inherently dynamic such that dispersal-mediated connections among populations are difficult to predict and vary by place and time (Cowen & Sponaugle, 2009; Gaggiotti, 2017; Liggins, Treml, & Riginos, 2013; Watson, Kendall, Seigel, & Mitarai, 2012). Yet, knowledge regarding the sources and destinations of dispersive larvae underpin fundamental ecological and evolutionary dynamics and inform optimal management of marine resources for fishing and the protection of key habitats (Beger et al., 2010; Gaines, White, Carr, & Palumbi, 2010; Hock et al., 2017; Krueck et al., 2017).

Coupled biological-physical models (Cowen & Sponaugle, 2009; Werner & Cowen, 2007) relying on simulations based on species attributes and physical oceanography are increasingly being used to predict spatial and temporal aspects of planktonic larval dispersal. The flexibility of both spatio-temporal scale and outputs makes biophysical models extremely well-suited for alignment against various sources of empirical dispersal data (Cowen & Sponaugle, 2009; Jones, 2015; Kool, Moilanen, & Treml, 2013; Liggins et al., 2013) and for generating detailed spatial predictions useful for management assessments (Hock et al., 2017; as in 2016; Krueck et al., 2017).

The extent to which biophysical models accurately approximate real dispersal phenomena, however, remains an open question. As models of open natural systems, strict validation or rejection of biophysical dispersal models is not feasible (Oreskes, Shrader-Frechette, & Belitz, 1994). Rather, such models can be evaluated against empirical data whereby alignment of the model and empirical data confers increased confidence that both the model and empirically-derived statistics describe the phenomenon of interest, in this case dispersal of planktonic larvae. Geographic surveys of intraspecific genetic variation can provide important insights regarding dispersal (Hellberg, 2009; Riginos & Liggins, 2013; Selkoe et al., 2016a) and indeed several studies have considered predictions arising from biophysical dispersal models alongside observed spatial genetic patterns, typically for coastal marine taxa. Although some individual studies report correlations (Benestan et al., 2016; Dalongeville et al., 2018; Foster et al., 2012; Galindo, Olson, & Palumbi, 2006; Schunter et al., 2011; Thomas et al., 2015; Truelove et al., 2016; White et al., 2010; Xuereb et al., 2018), a recent review of the field found only moderate to low concordance between biophysical predictions and empirical genetic patterns (Selkoe, Scribner, & Galindo, 2016b).

There are many reasons why biophysical dispersal models and genetic data may not align. First, biophysical models typically focus solely on dispersal and do not consider differential recruitment and/or reproductive success of settlers (Pineda, Hare, & Sponaugle, 2007). Second, a biophysical model might not sufficiently account for important biological attributes of larvae or complex near shore oceanography, which is notoriously difficult to parameterise (Metaxas & Saunders, 2009; Pineda et al., 2007; Werner & Cowen, 2007). Third, there is a mismatch in time scales, where biophysical models draw upon oceanographic information collected within recent years or decades whereas genetic inferences, especially those based on population allele frequency differences, arise from long term processes (thousands of years and longer). Fourth, population allele frequencies do not solely reflect gene flow resulting from dispersal but are also shaped by changes in population sizes, range expansions, colonization and so forth (Whitlock & McCauley, 1999). Lastly, although biophysical models typically predict asymmetric directionality of dispersal, typical descriptors of genetic diversity are symmetric so that statistical comparisons between biophysical models and empirical genetic data nearly exclusively rely on transforming modelled dispersal predictions into symmetric measures, greatly reducing the information content (Kool et al., 2013; Riginos, Crandall, Liggins, Bongaerts, & Treml, 2016).

Only a few studies have attempted to ascertain whether empirical genetic patterns are specifically consistent with the asymmetric dispersal processes inherent to biophysically derived dispersal predictions. For example, migration matrices derived from biophysical modelling were used to inform gene flow in forward population genetic simulations (Galindo et al., 2006) and to analytically predict genetic differentiation based on forward matrix projections (Kool, Paris, Andréfouët, & Cowen, 2010; Kool, Paris, Barber, & Cowen, 2011), yielding outcomes that qualitatively matched empirical genetic patterns. For the Caribbean coral *Orbicellea annularis*, genetic distances derived from a forward matrix project were shown to be well correlated (ρ = 0.49) to empirical genetic distance estimates (Foster et al., 2012). Another strategy draws upon historical demographic simulations to quantify migration values consistent with empirically observed allele frequency spectra or DNA sequences. For example, Matz *et al*. (2018) documented correlation between population pairwise estimates of migration derived from allele frequency spectra against biophysically derived migration probabilities for five populations for the coral *Acropora millepora* from the Great Barrier Reef (GBR) (Mantel R = 0.58, P = 0.05), with a preponderance of southward migration consistent between both measures. Comparing all sampled populations holistically, Crandall *et al*. (2012) constructed a series of coalescent models of gene flow and contrasted the likelihood of observed DNA haplotype distributions from the biophysically-informed gene flow models against various other null geographic models; they demonstrated greater likelihood of biophysically informed gene flow models across three species of nerite snails in the South Pacific. Recently, three studies turned to asymmetric eigenvector mapping (AEM: Blanchet, Legendre, Maranger, Monti, & Pepin, 2011) where asymmetric processes (such as biophysical migration probabilities) are statistically modelled as spatial autocorrelation structures; inclusion of AEMs substantially improved predictions of spatial genetic structure for American lobster (Benestan et al., 2016), California sea cucumbers (Xuereb et al., 2018), and Mediterranean striped red mullet (Dalongeville et al., 2018). Thus, these first few studies that quantitatively incorporate asymmetric biophysical predictions suggest that directions of larval dispersal are important elements of marine population connectivity.

Here, we comprehensively assess the alignment between biophysical models of dispersal and observed spatial genetic patterns for two common broadcast spawning coral species on the GBR, *Acropora tenuis* and *Acropora millepora*, drawing upon methods based on historical demographic simulations, patterns of shared alleles, and AEM spatial autocorrelation structures. We capitalise upon spatially rich genetic data sets for the two species (Lukoschek, Riginos, & van Oppen, 2016; van Oppen, Peplow, Kinnimonth, & Berkelmans, 2011) with sampling encompassing most of the 2300 km extent of the GBR. Oceanographic patterns suggest that directional gene flow is likely for GBR species such as *A. tenuis* and *A. millepora* (Fig. 1a).

**Figure 1:**
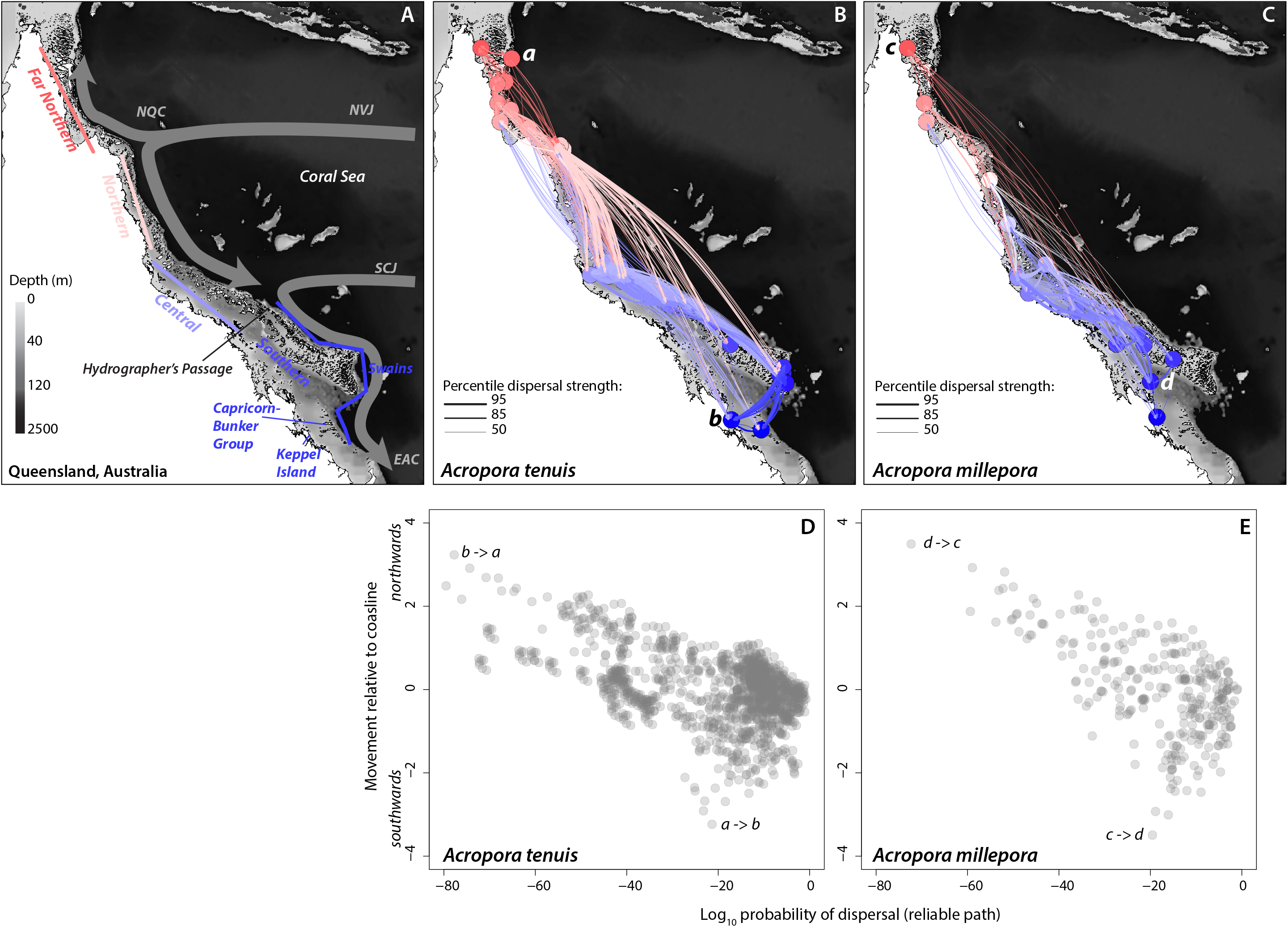
Sampling locations and main attributes of biophysical model. A) Coastal Queensland and the Great Barrier Reef where bathymetry is shown by grey shading and 120 m depth reflects the approximate land mass exposure at lowest Pleistocene sea level stands. Designated regions correspond to management areas. Major offshore currents are shown (NVJ and SVJ: north and south Vanuatu jets, NQC: north Queensland current, EAC: east Australia current; modified from Coukroun *etal*. 2010; Mao & Luick 2014). B & C) Summary of top 50 percentile predicted connections based on relative path probabilities for *Acropora tenuis* (B) and *Acropora millepora* (C). Sampling locations are color coded by latitude with northern low latitude sites shown in reds (warm) and southern higher latitude sites shown in blues (cool). Vectors show predicted dispersal probabilities with thicker lines indicating higher probabilities and colored by source population. D & E) Dispersal probabilities relative to coastline position for *Acropora tenuis* (D) and *Acropora millepora* (E). The largest positional changes northwards have the lowest probabilities whereas small positional changes and movements southwards have higher probabilities. Movement relative to coastline is based on the difference in relative position between locations as summarised by the first principal component axis describing Queensland coastline. Example contrasts in northward vs. southward dispersal strengths shown for example population pairs (a,b and c,d) traversing the length of the sampling region where a->b and c->d are high probabilities (P ¾ 10^−20^) of southward movement (−4 relative positional change) and b->a and d->c are low probabilities (P ¾ 10^−40^) of northward movement (+4 relative positional change).

Although present-day oceanography implies that larval dispersal can create connections among distant reefs, several species show differentiation between central/northern vs. southern GBR locations (reviewed by van Oppen et al., 2011), indicative of possible historical vicariance associated with past sea level changes. This divergence is most notable for the spiny chromis damselfish for which genetically distinct colour morphs abut at the north end of Hydrographer’s Channel (Planes & Doherty, 1997). Recent geological investigations, however, have identified reef structures along the Australian shelf edge that may have provided coral reef habitat during intermediate and low sea level stands (Abbey, Webster, & Beaman, 2011; Hinestrosa, Webster, & Beaman, 2016), implying that reef species may have shifted their range margins through multiple glacial cycles following the depth contours of available coastline but maintaining their approximate latitudinal positions.

No single approach can simultaneously infer modern and historical gene flow regimes across numerous sites spanning large spatial scales. Rather than discounting historical influences (as is implicit in many spatial genetic methods), we use coalescent demographic simulations to evaluate competing scenarios of long-standing gene flow vs. late Quaternary divergence. The resultant affirmation of gene flow-dominated demography along the GBR justifies subsequent frequentist analyses focusing on directional gene flow inferred from population allele frequencies and using all sampled reefs. Treating the biophysical models as hypotheses of spatial connections among GBR reefs for *Acropora* corals, we test whether projected asymmetric directions and dispersal strengths are superior predictors of spatial genetic patterns as compared to simple symmetric null predictors. In addition, we examine the spatial scales and regions for which biophysical predictors best align to observed genetic patterns. This study provides a framework for aligning spatially rich population genetic data against *a priori* predictions of asymmetric dispersal and represents the most comprehensive analysis of asymmetric gene flow along the full length of the GBR to date.

## METHODS

### Multi-step connectivity pathways based on larval dispersal models

Connectivity modelling follows protocols described in detail by Hock et al. (in prep) with an overview in the supplementary materials. The resultant species-specific connectivity networks (based on all 3806 GBR reefs) were used to predict multistep paths between the reefs from which the genetic samples were obtained, focusing on 1) the minimum number of *stepping-stones* between two reefs (Dijkstra’s 1959), 2) the *maximum flow* capacities of links necessary to connect the two nodes in a network (Ford & Fulkerson, 1956; Boykov & Kolmogorov, 2004), and 3) the *reliable path* represented by the maximum product of link weights representing the greatest chance of (direct or multi-step) larval exchange (Hock & Mumby, 2015). See supplemental methods and Sup. Fig. 1 for details and further explanation.

### Genetic data

We capitalise on the extensive microsatellite datasets for *Acropora tenuis* (Lukoschek et al., 2016) and *Acropora millepora* (van Oppen et al., 2011). Some populations were merged or omitted to match the larval dispersal model, some individuals with high missing data were excluded, and conformity to Hardy-Weinberg expectations and linkage equilibria were verified. See supplemental methods.

### Resolving historical influences using coalescent-based ABC

If allele frequencies among *Acropora* populations have been strongly influenced by past interruptions to gene flow, then we would need to evaluate observed geographic differentiation in light of such a divergence history. In contrast, if gene flow has dominated the modern distributions of allelic diversity then dispersal interpretations based on observed allele frequencies are reasonable. We used an approximate Bayesian computation (ABC) approach to test the competing hypotheses of historical divergence vs. stepping-stone gene flow, with the location of historical divergence set at the northern opening of Hydrographer’s Passage (Fig. 1). Owing to the high computational requirement in simulating genetic data to match our full datasets, we used representative populations for our ABC analyses (Fig. 2). A custom R script drew parameter values from the prior distributions, which were used with fastSimcoal2 v2.6.0.3 (Excoffier, Dupanloup, Huerta-Sánchez, Sousa, & Foll, 2013) to generate a total of 500,000 simulations, which were summarised using statistics calculated by Arlequin (arlsumstat v3.5.2.2: Excoffier & Lischer, 2010). Details of simulation conditions, model support, and cross-validation procedures are in the supplementary methods.

**Figure 2:**
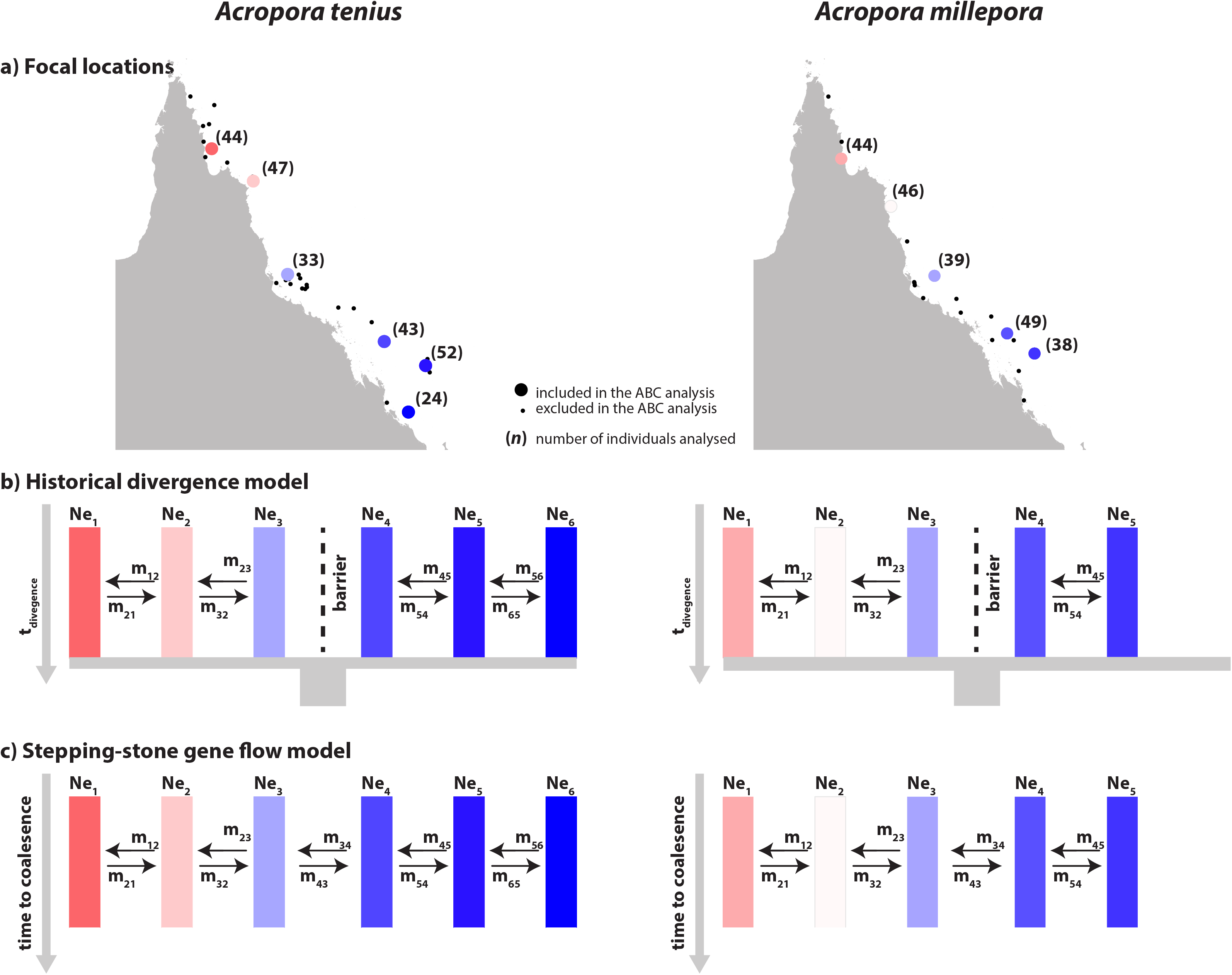
Competing models of historical scenarios evaluated using Approximate Bayesian Computation based on historical divergence vs. gene flow between adjacent stepping stone populations. A) Representative populations chosen for comparison via ABC against simulated data. B) Historical divergence models with a barrier to gene flow at Hydrographer’s Passage. C) Stepping-stone gene flow models where migration was permitted only between adjacent populations but with rates free to vary. The prior distributions of the parameters are summarized in Table S1. ABC analyses provided higher support for the stepping-stone gene flow models (A. *tenuis:* posterior probability = 0.999; Bayes Factor = 1210; *A. millepora:* PP = 0.647; BF = 3) compared to the model of historical divergence vicariance.

### Asymmetric gene flow and larval dispersal connectivity

To estimate directionality of gene flow we use information from semi private alleles between pairs of populations (Sundqvist, Keenan, Zackrisson, Prodöhl, & Kleinhans, 2016) as implemented by divMigrateOnline (https://popgen.shinyapps.io/divMigrate-online/) using the G_ST_ option (with no threshold for filtering out non-significant values). We assessed the correlation between these asymmetric (directional) estimates of gene flow with asymmetric measures of connectivity derived from the larval dispersal model (described above). Significances of the correlations were evaluated using 10,000 permutations. Maximum flow and reliable path metrics were (natural) log transformed for these and subsequent evaluations.

### Spatial eigenvector mapping

Although isolation-by-distance approaches have long been applied in spatial genetic contexts, they fail to account for spatial autocorrelation structures. Moran Eigenvector Maps (MEMs) model orthogonal spatial structures and, when combined with multivariate analyses, can test for spatial regression (Borcard & Legendre, 2002; Dray, Legendre, & Peres-Neto, 2006) and are well-suited for genetic investigations (Diniz-Filho, Nabout, Telles, Soares, & Rangel, 2009). Asymmetric eigenvector maps (AEMs) extend spatial modelling to describe spatial structures arising from directional processes such as ocean currents (Blanchet et al., 2011; Blanchet, Legendre, & Borcard, 2008). Here we use AEMs derived from the biophysical model to test whether asymmetric (directional) connectivity distances predict genetic structure, with AEM model fits contrasted to null symmetric expectations described by MEMs. Creation of link structures, weighting schemes, and connexion diagrams (Blanchet et al., 2008; 2011) are detailed in the supplemental methods and Supp. Fig. 1. Observed population allele counts were chi-square transformed (somewhat upweighting rare alleles: Legendre & Gallagher, 2001) to form the response variables for MEM and AEM analyses. Selection of MEM and AEM model was undertaken with forward model selection based on adjusted R^2^ values following Dray *et al*. 2006 and Blanchet *et al*. 2011. Significance values of individual AEMs for the final model were determined using redundancy analysis.

## RESULTS

### Larval dispersal model

Larval dispersal models for *Acropora tenuis* and *Acropora millepora* suggest that GBR populations are well connected when multi-step (i.e. multigenerational) connections are considered. Overall southward connections were more prevalent, especially long-distance connections: this can be visualised by vectored arcs curving the right in Fig. 1B and C along with greater dispersal probabilities for southwards movements (Fig. 1 D and E). However, some strong northward connections were also present (vectored arcs curving the left) especially among central reefs and some central to northern reefs. All three multistep connectivity metrics considered were highly correlated (ρ ≥ 0.86 for *A. tenuis;* ρ ≥ 0.80 for *A. millepora)* with the greatest correlations between stepping stone distance and reliable paths (ρ ≥ 0.98 for both species). Given these high correlations, we present results primarily for the reliable path metric, which arguably best aligns with biological intuitions regarding population connectivity across time (Hock & Mumby, 2015).

### Resolving historical influences using coalescent-based ABC

In the present study, it was necessary to verify that gene flow has shaped *Acropora* spatial genetic patterns and exclude vicariance as an alternative scenario before proceeding with analyses that implicitly ignore divergence. Indeed, ABC analyses yielded higher support for the gene flow only stepping-stone model (A. *tenuis:* posterior probability, PP= 0.999; Bayes Factor, BF = 1210; *A. millepora:* PP= 0.647; BF = 3) compared to a model of historical divergence vicariance. Applying model selection with the pseudo-observed data (POD) yielded high accuracy (> 99; proportion of POD that were correctly supported) and high robustness (> 99 with thresholds above 0.6 PP). Overall, these results indicate strong support (sensu Roux et al., 2016) for the gene flow only stepping-stone model for both species.

### Asymmetric gene flow and larval dispersal connectivity

For *A. tenuis*, all directional predictors of connectivity yielded significant correlations with relative gene flow estimates from divMigrate (ρ values for stepping stone distance: −0.39; maximum flow: −0.31; reliable path: −0.37; in all cases P << 0.001). For *A. millepora*, reliable path (ρ = −0.16; P < 0.005) and stepping stone distances (ρ = −0.13; P < 0.03) were significantly correlated with relative gene flow but maximum flow was not (ρ = −0.06; NS). For both species, inspecting the relationship between reliable path probabilities and gene flow (Fig. 3) shows that for populations predicted to be well connected by the larval dispersal model (i.e. P ≥ 10^−20^) estimates of relative gene flow were highly variable (~0-1), whereas sites predicted to require many stepping stone connections (P ≤ 10^−40^) had consistently lower gene flow estimates. This pattern was stronger in *A. tenuis* than *A. millepora*.

**Figure 3:**
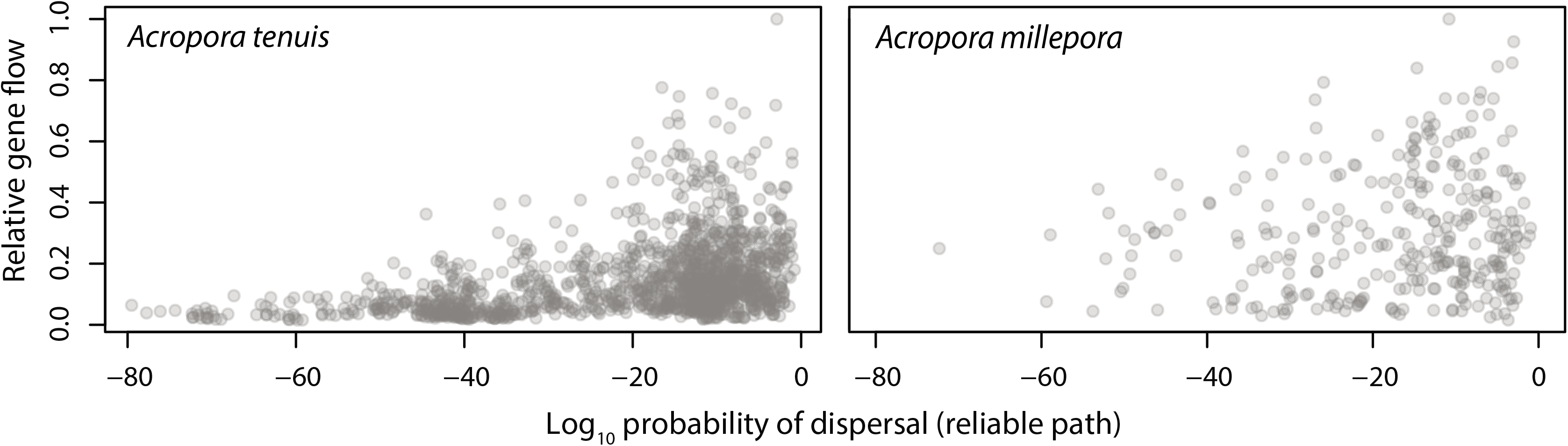
Concordance in directional movement between the larval dispersal model and genetic estimates for pairs of populations for *Acropora tenuis* (A), and *Acropora millepora* (B). Routes of low predicted connectivity (small relative path probabilities: P < 10^−40^) experience low directional gene flow (using the DivMigrate method). Routes of high predicted connectivity (P > 10^−20^) have variable rates of gene flow.

### Spatial eigenvector mapping

Directional spatial autocorrelation as assessed by AEMs explained a greater proportion of variance in allele frequencies for both species (Table 1), where the best models were based on alongshore north to south movements with inshore connections from the Swains to Keppel Island and to the Capricorn-Bunker group (for *A. tenuis*, R^2^ = 0.37 with 9 AEMs retained, and for *A. millepora*, R^2^ = 0.27 with 5 AEMs retained). There was no consistency in terms of best-performing link weighting (i.e. stepping stone connections, maximum flow, and reliable path) and notably the binary weighting (presence vs. absence, no weighting by distance) for north to south connections in *A. millepora* was the best performing, indicating that simply recognising strong connections (i.e. those with reliable path probabilities in the top 50^th^ percentile) yields good approximations of spatial genetic patterns. Fig. 4 depicts the highest scoring AEMs for *A. tenuis* where AEM 1 (3.5% of allele frequency variance; P = 0.001) as the largest scale of positive autocorrelation showed a gradient across the entire sampled region of the GBR and AEM 5 (2.3 % of variance; P = 0.001) described more local scale autocorrelation patterns. For *A. tenuis*, AEMs 1, 5, 17, 3, 2, and 12 were individually significant at a P = 0.05 threshold based on an RDA of the final model where higher numbers (i.e. AEM17) indicate finer spatial grain sizes out of a total of 19 possible AEMs of positive spatial autocorrelation. For distance-based *A. tenuis* MEM models, the best model (PCNM) retained MEMs 4, 15, 35, 22 indicating a mix of large- and fine-scale autocorrelations. Similar results were found for *A. millepora* (Supp Fig. 3) with AEM 1 describing the greatest amount of spatial variance in allele frequencies (5.9%, P = 0.003) followed by AEM 8 (5.7 %, P = 0.007), and AEMs 1, 8, 5, and 3 individually significant below a P = 0.05 threshold out of 9 AEMs evaluated. For distance-based *A. millepora* MEM models, the best model (custom saturated) retained only MEM 12. For *A. tenuis*, sampling within regions was sufficiently dense to evaluate fits for adjacent regions of competing AEM models. In all instances links based on north to south alongshore flow returned the best scoring models, with fit to allelic frequency patterns notably lower in the far northern and northern reefs (Table 2).

**Figure 4:**
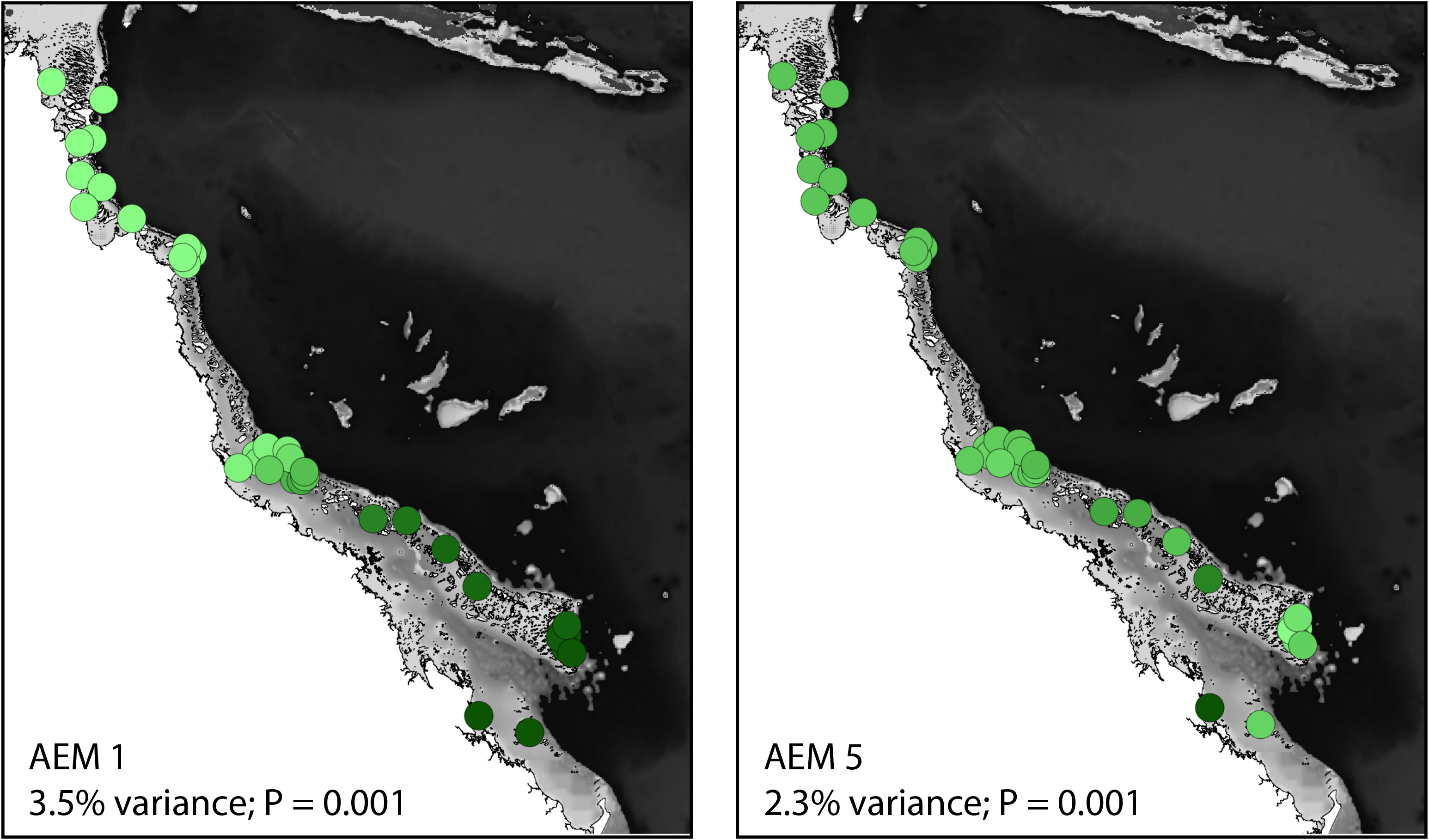
Leading asymmetric eigenvector maps describing spatial genetic structure in *Acropora tenuis*. Values by sampling location are colored by intensity of green hue such that locations with similar colors have similar AEM values. The greatest proportion of variance (AEM 1) describes a GBR-wide cline in genetic diversity followed by local scale spatial autocorrelation structures (AEM 5).

**Table 1:** Spatial eigenvector map modelling of genetic structure. The models explaining the greatest amount of genetic variance are those that incorporate directional link structures derived from the larval dispersal model, especially those that are based on north to south alongshore flow.

| Eigenmodel | Link structure | *Acropora tenuis* |  |  | *Acropora millepora* |  |  |
| --- | --- | --- | --- | --- | --- | --- | --- |
|  |  | Link weighting of best model <sup>a</sup> | Numb EMs in best model <sup>b</sup> | adjusted R<sup>2</sup> <sup>b</sup> | Link weighting of best model <sup>a</sup> | Numb EMs in best model <sup>b</sup> | adjusted R<sup>2</sup> <sup>b</sup> |
| MEM | Delaunay | Binary | 1 | 0.02 | Binary | 1 | 0.02 |
| MEM | Delaunay | log | 1 | 0.03 | log | 1 | 0.03 |
| MEM | PCNM | y=1 | 4 | 0.24* | log | 1 | 0.09 |
| MEM | Custom saturated | y=1 | 3 | 0.16* | log | 1 | 0.10 |
| AEM | Alongshore N to S only <sup>c</sup> | MF, y=1 | 9 | 0.37 | Binary | 5 | 0.27 |
| AEM | Alongshore both N to S and S to N | SS, y=1 | 8 | 0.29 | MF, y=1 | 2 | 0.15 |

<sup>a</sup> Binary distances represent a single model. The link structures were evaluated for link distance weightings based on the formula $1 - (D/D_{max})^y$ for $y=[1-3]$ or with a log transformation to approximate a typical isolation-by-distance analysis. For AEMs, links were limited to those in the top 50 percentile of reliable path weights. Link weightings were evaluated based on the number of stepping stone (SS) connections, maximum flow (MF), and reliable path (RP) strength.
<sup>b</sup> Following forward model selection; asterisks indicate spatial models with $\Delta AICc < 2$ as compared to the null (intercept only) model.
<sup>c</sup> Select AEMs shown graphically in Fig. 4 and Sup. Fig. 3

**Table 2:** Asymmetric eigenvector map modelling regions within *Acropora tenuis*

| Region | Link structure | Best model by region |  |  |
| --- | --- | --- | --- | --- |
|  |  | Link weighting<br>*a* | Numb EMs in<br>best model *b* | adjusted R<sup>2</sup> *b* |
| Far north & north | Alongshore both N to S<br>and S to N | MF, y=1 | 3 | 0.17 |
| North, central, south | Alongshore N to S | MF, y=1 | 7 | 0.32 |
| Central & south | Alongshore N to S | RP, y=2 | 5 | 0.35 |

<sup>a</sup> Link structures were evaluated for distance weightings based on the formula $1-(D/D_{\max})^y$ for y=[1-3].
<sup>b</sup> Following forward model selection.

## DISCUSSION

Asymmetric dispersal is undoubtedly a common attribute of benthic marine species. For the corals *Acropora tenuis* and *Acropora millepora*, a preponderance of north to south movement along the GBR was predicted by biophysical models and this directional signal was matched in patterns of microsatellite allele frequencies. Although previous studies have noted distinct characteristics of northern + central vs. southern + Swains + Capricorn-Bunker regions (Lukoschek et al., 2016; van Oppen et al., 2011), here we can confirm asymmetric gene flow as a contributor to these regional differences. Alternative scenarios involving historical divergence and symmetric gene flow were poor matches to observed spatial genetic patterns relative to scenarios based on directions and extent of larval exchange predicted by biophysical models. Thus, at least at the regional scale within the GBR, oceanography is likely to strongly influence connections arising from planktonic larval dispersal.

### Regional connections among GBR populations

Whereas geographically restricted alleles are a hallmark of long-standing isolation, highly directional dispersal such as among coastal populations can result in regionally private alleles in downstream populations if dispersal is sufficiently rare and when accompanied by high self-recruitment of settlers to their parental populations (Pringle & Wares, 2007). This expectation qualitatively matched diversity gradients for both *Acropora* species (as noted by Lukoschek et al., 2016). Here, coalescent demographic simulations allowed us to formally evaluate a hypothesis of Quaternary divergence against a scenario of long-standing gene flow without divergence. For *A. tenuis*, we found very high support for gene flow without divergence (posterior probability: 0.999; BF = 1210) and moderate support for this scenario in *A. millepora* (PP= 0.647; BF = 3). Thus, if there were past divergence events, the imprint of such events is no longer discernible from extant allele frequencies and present-day population genetic structure should be strongly influenced by recent gene flow.

Indeed, contemporary biophysical expectations for gene flow were strong predictors of genetic patterns especially at large spatial scales. Using shared alleles, the weakest genetically connected population pairs (negligible gene flow from reef *i* to reef *j*) match pairs having the weakest predicted multigenerational connection links (Fig. 3). Where the biophysical dispersal models predicted stronger connections, in contrast, inferred gene flow was highly variable indicating that at shorter distances the biophysical models and gene flow were not consistently aligned. Formal statistical testing for spatial structure of allelic distributions with MEMs and AEMs reinforced these findings, where eigenvectors derived from predicted asymmetric connections explained considerably more variance than eigenvectors derived from null geographic models of symmetric connections (Table 1). Additionally, for both *A. tenuis* and *A. millepora* AEM 1, which describes the largest scale pattern of spatial relationships (Fig. 4 and Sup. Fig. 3), explained the greatest proportion of variance in allelic distributions (3.5 and 5.9%, respectively). AEM analyses, however, were also consistent with spatial patterning at smaller spatial scales arising from asymmetric connections (Table 1).

That biophysical models and spatial genetic structure were misaligned at small scales does not seem overly surprising. Small-scale oceanography is likely to be highly variable and may lead to higher levels of mixing among local clusters of reefs than a limited number of dispersal years would suggest. Moreover, the time periods are incongruent as the dispersal models spanned four years (2008-2012), whereas genetic data were derived from mature colonies (collected in 2009-2013 for *A. tenuis* and 2002-2009 for *A. millepora)*. Additionally, chaotic genetic patterns at small scales such as those observed here (Fig. 3) are well-known for marine taxa, perhaps reflecting stochasticity associated with planktonic dispersal and/or post-settlement selection (e.g. Johnson & Black, 1984). Finally, biophysical larval dispersal models implicitly assume that post-settlement mortality rates would not be affected by origins of settlers, but larval origins could influence juvenile fitness (Marshall et al. 2010).

Although spatial distributions of microsatellite alleles for both *A. tenuis* and *A. millepora* were broadly consistent with asymmetric dispersal predicted by biophysical models, the best alignments were obtained only considering the approximately north to south connections along the length of the GBR (Table 1). Notably biophysical models for both species suggested substantial counterflows (south to north) especially among inshore reefs (Fig. 1), yet adding these connections to AEM models did not increase model fit (Table 1). Thus, either the stronger north to south connections are sufficient for capturing most of the microsatellite allelic variance or the predicted south to north movements are not realised: for example, larvae may conceivably disperse northwards but could have low fitness and therefore not make substantive contributions to gene flow. Matz *et al*. (2018) also detected greater southwards migration among five populations of GBR *A. millepora* based on ~11,500 nucleotide polymorphisms, albeit with some northwards migration and only considering an inshore southern reef location (Keppel Island). Southward spread also characterises Crown-of-Thorns starfish outbreaks that appear to originate in northern reefs (Pratchett et al., 2014). Therefore, our finding of a strong southward dispersal signal is compatible with these other results, but the sensitivity of AEM based analyses to bidirectional flows is unclear.

For *A. tenuis*, we were able to further assess correlations between biophysical predictions and allelic spatial distributions between adjacent regions (Table 2). For southern and central reefs, north to south connections models yielded the best fits to the data (R^2^ ≥ 0.32). These observations spatially align with major jets from the Coral Sea encountering the GBR and driving southward advective flows along the GBR (i.e. surface waters of the South Caledonia Jet at the top of the central region and Eastern Australian Current at the Swains: Choukroun, Ridd, Brinkman, & McKinna, 2010; Mao & Luick, 2014). In contrast, far northern and northern reefs showed reduced concordance between predictions and genetic observations (R^2^ = 0.17) with a model based on bidirectional connections yielding the best fit to this regional data; surface waters of the North Caledonia Jet encounter the GBR here and creates both northward and southward flows and thus might explain a possible signal of bidirectional connections among *A. tenuis* populations (Choukroun et al., 2010; Mao & Luick, 2014). For both species, biophysical models predict rare dispersal connections especially among far northern reefs (Fig. 1) and yet negligible population structure was observed for either *A. tenuis* or *A. millepora* among reefs in the far north (Lukoschek et al., 2016) suggesting a weaker fit between dispersal models and observed genetic patterns in this region.

In summary, for the two acroporid corals considered here, we find strong support for recent gene flow in a predominantly north to south direction. This result implies that northern and central reefs are important sources for downstream southern reefs, at least over evolutionary time scales of many generations. Southern outer shelf reef populations, however, harbour greater genetic diversity than northern reefs (Lukoschek et al., 2016) and are therefore both valuable in terms of distinctive diversity and, as the present results show, likely to be strongly self-recruiting so that they may be more self-sustaining than central and northern reefs. Dynamics of inshore reefs may be different from outer shelf reefs, for example, Lukoschek et al. (2016) highlighted the genetic distinctiveness of the inshore Keppel Island population for both *A. tenuis* and *A. millepora* and Matz et. al. (2018) found low genetic diversity in the Keppels but with no sampling of southern outer reef populations as a basis for comparison. Matz et. al. (2018) also reported low-levels of northward gene flow among their inner reef populations (e.g. Keppel to Magnetic and Orpheus Islands). Further genomic studies with more comprehensive spatial sampling may yield greater detailed resolution regarding rates and directions of gene flow, as the microsatellite data we examined here lack the power to confidently estimate specific gene flow rates.

With temperatures and extreme heating events projected to increase in frequency (Wolff et al., 2018), resolving the GBR-wide capacity for gene flow including the directionality of naturally occurring gene flow provides information relevant to discussions regarding assisted migration and genetic rescue (Anthony et al., 2017). For example, although the southern outer reefs, especially the Swains, have been largely spared from bleaching events, our results indicate that naturally occurring dispersal from south to north is most likely ecologically insignificant (Fig. 3), and thus would indicate that prospective migrants from the southern regions would be unlikely to recolonise central and northern reefs in sufficient quantities over decadal time scales. Because naturally occurring genetic rescue relies on evolutionarily significant (and not ecologically significant) gene flow, the prospects for this phenomenon are better, although predicated on receiving locations hosting healthy populations. Should adaptive variants (such as conferring heat tolerance or bleaching resistance) exist in some reefs then they might be able to spread naturally, with southward spread more probable. In any event, increased knowledge regarding the directionality and magnitudes of genetic exchange among GBR reefs will help forecast possible future evolutionary dynamics and can help identify dispersal barriers that could be circumvented via assisted migration (Hoffmann et al. 2015). In the case of GBR corals, assisted movements in a northward direction would bypass natural dispersal limitations.

### Detecting contemporary asymmetric planktonic larval dispersal

Relatively few studies have evaluated the effect of asymmetric larval dispersal on spatial genetic patterns in marine species (Riginos et al., 2016). Perhaps not surprisingly, these studies, like our results here with *A. tenuis* and *A. millepora*, consistently uncover substantial asymmetries in inferred dispersal (e.g., Benestan et al., 2016; Dalongeville et al., 2018; Xuereb et al., 2018). Clearly, conventional analyses, especially those based on summary statistics like F_ST_ where all directional information is obscured, will fail to uncover important elements regarding relationships among populations (Kool et al., 2013; Riginos et al., 2016).

An additional challenge is to clarify the time frame for processes yielding asymmetric spatial genetic patterns. Empirical genomic data paired with specific historical demographic simulations can sometimes put bounds on the timing and directionality of gene flow (as in Duranton et al., 2018; Matz et al., 2018), but such approaches are largely constrained to examining a few populations at a time due to exponential increase in number of potential free parameters (for example, if migration is free to vary among n populations then there will be n!/(n-2)! parameters for directional migration). Moreover, there is greater uncertainty surrounding parameter estimation for recent events (Robinson, Coffman, Hickerson, & Gutenkunst, 2014). Simple frequentist approaches based on linear models such as AEMs represent an attractive alternative especially as they have been conceived for uncovering contemporary structuring processes in species composition of ecological communities (Dray et al., 2006), but if applied to intraspecific genetic data these approaches are likely to be misleading when historical events have strongly shaped spatial genetic patterns.

In the present study, we attempted to bridge this historical-contemporary dilemma by using constrained historical demographic simulations to reject the historical scenario most likely to influence observable spatial genetic patterns for GBR species, giving us greater confidence that the spatial genetic patterns we observe for *A. tenuis* and (to a lesser extent) *A. millepora* reflect evolutionary processes from the recent past. Using prior information based on geology, historical habitat shifts, or histories of co-distributed species to identify relevant alternative hypotheses and then evaluating those hypotheses against hypotheses based on contemporary gene flow (such as simple stepping-stone gene flow as used here) is a reasonable check of data before proceeding to interpret results from linear models (including AEMs) at face value.

### Biophysical models as summaries of larval dispersal

Biophysical models are increasingly being used to make detailed spatial predictions of planktonic larval dispersal often with the aim of guiding management actions (as in Hock et al., 2016; 2017; 2019; Krueck et al., 2017). Although such complex models of natural systems cannot be truly tested (Oreskes et al., 1994), alignment of model outputs against independent biological data can provide greater confidence that a biophysical larval dispersal model captures elements of biological reality. In the present study, we find that directions and relative magnitudes of dispersal connections derived from biophysical models are better predictors of spatial genetic structure in *A. tenuis* and *A. millepora* than null models of symmetric relationships, providing some confirmation that real biological processes mirror expectations arising from biophysical simulations. The superior predictive power of asymmetric predictors provides further evidence of the need to consider directional flow when analysing connectivity in marine systems.

For our *Acropora* species on the GBR, the strongest alignments between the models and empirical genetic data are at large spatial scales (among regions) and involve north to south connections. Because small numbers of migrants can homogenize allele frequencies between populations, inferences based on allele frequencies are poorly suited to distinguishing strengths of demographically significant connections (Waples, 1998). Allele frequency based methods, however, can theoretically discriminate between rare and very rare connections such as those that are likely to link geographically distant populations (including via multigenerational connections). Thus, the appropriate spatial scale for evaluating biophysical larval dispersal models with population genetic data is at large spatial distances such a regional connections among GBR coral populations where we find the greatest concordance between predicted and observed patterns. Comprehensive evaluation of biophysical models will best be undertaken with a variety of complementary empirical data types (Gaggiotti, 2017; Jones, 2015), where geographic patterns of allelic distributions contribute valuable information about rare long-distance connections. Here we have shown that the directionality of such rare long-distance connections discerned from spatial distributions of alleles provides additional useful information for gauging concordance of larval dispersal models against past gene flow.

## Supporting information

Supplementary methods and results

## Acknowledgements

We thank FG Blanchet for guidance on creating connexion diagrams and C Doropoulos for help with parameterisation of the dispersal models, the QRIS computing cluster for enabling simulation modelling, and funding support from the Great Barrier Reef Foundation.

## Data accessibility statement

Raw data available at datadryad.org/resource/doi:10.5061/ dryad.h8gh3 for *Acropora tenuis* and datadryad.org/resource/doi:10.5061/dryad.q0834p71 for *Acropora millepora*. Scripts, datafiles, and example results are available on GitHub: github.com/khock1/CoralPaths; github.com/dinmatias; github.com/CRiginos1.

## Biosketch

The authors are linked by a desire to better understand how planktonic larval dispersal connects marine populations. Advances in measuring and predicting dispersal connections are urgently needed in the context of the Great Barrier Reef where there have been recent widespread outbreaks of coral bleaching. Thus, progress in uncovering the dynamics of long-distance dispersal connections contributes to an ultimate goal of identifying locations best-able to supply propagules and seed downstream reefs.

## Author contributions

CR and VL conceived the overarching concept and KH, AMM, PJM, & MJHvO contributed ideas fundamental to the study; VL, MJHvO, and KH collected the data; CR, KH, and AMM analysed the data and developed the figures; and CR led the writing with contributions from KH, AMM, PJM, MJHvO & VL.

## Notes

#### Summary of Updates

Some additional elaboration added to the discussion. Further explanation of methods appears in supplemental materials.

