## Supplementary methods and results for "Asymmetric dispersal is a critical element of concordance between biophysical dispersal models and spatial genetic structure in Great Barrier Reef corals"

*Multi-step connectivity pathways based on larval dispersal models*

Larval particles were released from 3806 indicative GBR reef polygons (GBRMPA 2004) and then displaced in 3D (to incorporate depth mixing and upwelling) using vectors representing hydrodynamic forces. The larval dispersal simulations were performed using CSIRO's Connie model for the GBR domain (www. csiro.au/connie), covering the continental shelf from Moreton Bay in Queensland to mainland New Guinea with a 4km horizontal spatial resolution for the forcing grid and a temporal resolution of one hour (Condie & Hepburn, 2012). Simulations were performed over 4 seasons (GBR summers 2008-2012), with realistic spawning dates for *Acropora* corals (using lunar phases and multiple spawnings per season if warranted). The particles followed general *Acropora* spawning behavior, with larvae released over the course of one night (or two nights on consecutive lunar cycles if there was split spawning) in a mass spawning event timed to match field observations (Babcock et al. 1986). A total of $10^4$ particles were released per spawning season from each of the indicative reef polygons. Particles were considered negatively buoyant (Arai et al. 1993), and dispersed in the surface layer of the oceanographic model domain. Once released, the particles were displaced in hourly intervals, and their distance to the reefs estimated at six hourly intervals. Larval particles could spend a maximum of 120 days post-spawning in the water column. Larval competency and survival functions were based on laboratory cultures of *Acropora* larvae (Connolly & Baird, 2010), providing time-senstive estimates of larval settlement upon their arrival at the destination reef. Refining the connections between reefs with time-sensitive mortality and competency to simulated larval particles (Connolly & Baird, 2010) allowed for assigning weighted (i.e. strength of exchange) asymmetric connectivity links between each source-destination pair of reefs to represent cumulative exchange over the simulated dispersal period.

The resultant region-wide species-specific connectivity networks (incorporating all 3806 GBR reefs) were used to predict multistep paths between the reefs from which the genetic samples were obtained. To calculate the minimum number of *stepping-stones* between two reefs in a connectivity network, all the positive link weights were set to 1. Links in this unweighted network include connections that were present at least once over the four years of simulations regardless of the strength of connection. Dijkstra's (1959) algorithm yielded the directed path with the minimum number of stepping-stones between any two reefs. *Maximum flow* metric calculations were performed by applying a maximum-flow minimum-cut algorithm (Ford & Fulkerson, 1956) on a weighted connectivity network in which link weights were considered to represent flow capacities. Determining the capacities of links necessary to connect the two nodes in a network (including multistep paths) then makes it possible to determine the maximum amount of flow between them. The maximum flow metric was calculated using the Matlab implementation of Boykov-Kolmogorov max-flow algorithm (Boykov & Kolmogorov, 2004). It is also possible to consider that link weights represent probabilities of dispersal (i.e., the probability that larvae from reef *i* will disperse to reef *j*). With links as dispersal probabilities, the probability of dispersal between a pair of sites is obtained by multiplying the link probabilities of all intermediate connections between them (Hock & Mumby, 2015). The most *reliable path* is then a pathway with the largest product of link weights representing the greatest chance of (direct or multi-step) larval exchange (for method of calculation, see Hock & Mumby, 2015). Supplemental Figure 1 visually depicts the relationships among these metrics.

*Genetic data*

The microsatellite datasets for *Acropora tenuis* (Lukoschek et al., 2016) and *Acropora millepora* (van Oppen et al., 2011) were filtered to exclude repeat multilocus genotypes per sampling location and to exclude individuals missing data for three or more loci. To align the genetic data against the larval dispersal model, we excluded three locations (22_SN_PIT, 23_HIGH_RK, 68_MAG_CAY) and merged some sites within reefs or islands as follows: Yonge (sites 10+11); No Name (sites 12+13); Palm Islands (sites 38-41,44); Heron and One Tree Island (sites 60-65) for *Acropora tenuis*. For *Acropora millepora,* we merged Sudbury 1 with Sudbury 2 and removed one locus (Am2_002) due to high missing data. In total, there were 10 loci for *A. tenuis* and 10 loci retained for *A. millepora*. Exact details of sampling locations and final samples sizes can be found in Supplementary Figure 2 and Supplementary Tables 2-3.

Using the R packages adegenet (ver. 2.0.1: Jombart & Ahmed, 2011) and pegas (ver. 0.9: Paradis, 2010) we tested for deviations from Hardy-Weinberg equilibrium and linkage equilibrium (based on the Zaykin, Pudovkin, & Weir, 2008 method for unphased multiallelic data). Since only a few combinations were significant below a 0.05 threshold (AT: 4 out of 390 locus-by-population combinations for HWE; 6 out of 1755 locus-by-allele-by-population combinations for LD; AM: 4 out of 209 for HWE, 6 out of 1045 for LD) we retained all loci that had passed the missing data filtering. For *A. tenuis*, we retained 1928 individuals from 39 sites and for *A. millepora*, we retained 917 individuals from 19 sites.

*Resolving historical influences using coalescent-based ABC*

Representative populations were used for our ABC analyses (Fig. 2) to make simulations computationally tractable. Parameter values were drawn from the prior distributions using a custom R script. These prior values were input to fastSimcoal2 v2.6.0.3 (Excoffier, Dupanloup, Huerta-Sánchez, Sousa, & Foll, 2013) to generate a total of 500,000 simulations of ten microsatellite loci for each genetic model. The simulated genetic data were summarised using the following statistics calculated with the Arlequin command-line version arlsumstat v3.5.2.2 (Excoffier & Lischer, 2010): average number of alleles, standard deviation of number of alleles, allele size range per population, and average pairwise $F_{ST}$ between populations. We limited the summary statistics to these measures to avoid extreme high dimensionality of summary statistics while retaining pertinent information about genetic diversity in each population and genetic differentiation between populations. In determining the support for each genetic model (model selection), we first obtained posterior samples from all the simulated data for each species by applying a threshold of 0.01, equivalent to the 10,000 simulated datasets closest to the observed genetic data. We then evaluated the support for each genetic model by performing a categorical regression (Beaumont, 2008) on the model indices and summary statistics of the posterior samples.

Following ABC model selection, we performed a cross-validation to verify that our approach could discriminate between historical divergence and stepping-stone gene flow scenarios and that the support for a model is not due to random chance. For the cross validation, we simulated an additional 10,000 data points for each model, to serve as pseudo-observed data (PODs). We then examined if the POD were correctly supported using a similar model selection procedure to that described in the main methods. Using the cross-validation results, we examined the minimum classification threshold for the posterior probability that will give a robustness of 0.95 (Roux et al., 2016). Empirical posterior probabilities greater than this threshold are deemed significant.

All the procedures for ABC model selection and validation were implemented using the R packages abc (ver. 2.1: Csilléry, François, & Blum, 2012) and nnet (ver: v7.3.12: Venables & Ripley, 2013). Posterior probabilities of each model were obtained from 20 trained neural networks with 15 hidden layers while weighing the posterior samples by Epanechnikov kernel leading to weights that are inversely proportional with the distance from the observed genetic data.

*Spatial eigenvector mapping*

Orthogonal spatial structures were modelled with Moran Eigenvector Maps (MEMs: Borcard & Legendre, 2002; Dray, Legendre, & Peres-Neto, 2006) and asymmetric eigenvector maps (AEMs: Blanchet et al., 2011; Blanchet, Legendre, & Borcard, 2008). For (symmetrical) MEMs, we followed Dray *et al.* (2006) and examined binary Delaunay, weighted Delaunay, and truncated (principal coordinates of neighbour matrices: PCNM) networks based on spatial position (i.e. geographic distance between sites) with a variety of link weighting schemes to alter the relative influence of short vs. long distance connections: all networks (excluding the binary Delaunay) were weighted by between site geographic distances for the functions 1-($D_{ij}$/Dmax)^y where $D_{ij}$ is the geographic distance between sites *i* and *j* and y =[1, 2, 3] (Dray et al., 2006). We also used the function ln(1-( $D_{ij}$ /Dmax)) to mirror conventional isolation-by-distance analyses. A custom saturated network allowing connections among combinations of sites was also used as an analogue to conventional isolation-by-distance analyses.

The construction of AEMs requires specifying connexion diagrams (Blanchet et al., 2008; 2011) indicating directionality of predicted process (in this case larval dispersal). Initial examination of connections for both species indicated a preponderance of strong connections (Fig. 1) southwards alongshore and westward cross-shelf (approximately NE to SW) especially connecting the Swains to Keppel Island and to the Capricorn-Bunker group. Several northwards alongshore connections were also predicted especially from outer shelf to central reefs. For AEMs, we constructed a series of connexion diagrams summarising the main system-wide connection directions following the procedure detailed in Blanchet *et al.* (2011), allowing north to south alongshore connections with an inshore bend in the southern GBR connecting the Swains to the Capricorn-Bunker group and Keppel Island. We also created connexion diagrams for south to north. In each case, models were fit with north to south only and in both directions. Connection links were created for all population pairs with reliable path probabilities in the top 50$^{th}$ percentile. Link weightings based on the number of stepping stone connections, maximum flow, and reliable path were evaluated. For *A. tenuis* where spatial coverage was more extensive, we also fit AEMs to adjacent sets of regions (far north and north; north, central and south; central and south).

Spatial eigenvector mapping analyses made use of R packages ade4 (ver. 3.3.2: Dray, Dufour, & Chessel, 2007), adespatial (ver. 0.0-8: Dray et al., 2017.), AEM (ver. 0.6: Blanchet, Legendre, & Gauthier, 2015), spdep (ver. 0.6-8: Bivand & Piras, 2015), and vegan (ver. 2.4-2: Oksanen et al., 2017).

**Supplementary figures:**

**Supplementary Figure 1:** Link structures and weighing schemes for spatial eigenvector models shown for an idealized set of sampling locations (black circles). Links are shown as grey lines. A) Moran Eigenvector Maps (MEMs) link structures depend solely on geographic position, where Delaunay triangulation and truncated criteria are commonly used to simplify the number of connections; connections between locations are bidirectional. B) MEM link weightings representing the strength of connections between locations can be unweighted (binary) or weighted where the increased influence of proximate locations is typically scaled by power functions (raised to the power of 1-3), and we also include a negative log scaling. C) Asymmetric Eigenvector Maps (AEMs) links are directional and indicated by arrowheads representing a directional process from top to bottom of the figure. Because the guidelines for constructing link structures for AEMs when sampling locations are irregular are not well established, we opted to simplify a saturated connection structure by only considering the more probable links using the reliable path criterion. D) AEM link weightings were based on three metrics: stepping-stone distance, maximum flow, and most reliable path probability. Considering two hypothetical locations S and T, the stepping-stone distance from S to T is the connecting path with minimum number of links, in this example a path of two steps shown in pink. The maximum flow is the sum of flow capacities through the minimum set of weakest connecting links that, if removed, would disconnect S from T, and the most reliable path is the product of link probabilities for the highest probability multistep pathway (shown in blue).

**Supplementary Figure 2**: Sampling locations for *Acropora tenuis* and *Acropora millepora*. Figures are modified from Lukoschek et al., 2016. Grey points indicate locations that were omitted for the present study and locations that were merged are shown in matching colours.

**Supplementary Figure 3:** Leading asymmetric eigenvector maps describing spatial genetic structure in *Acropora millepora*. Values by sampling location are colored by intensity of green hue. The greatest proportion of variance (AEM 1) describes a GBRF wide cline in genetic diversity followed by local scale spatial autocorrelation structures (AEM 8).

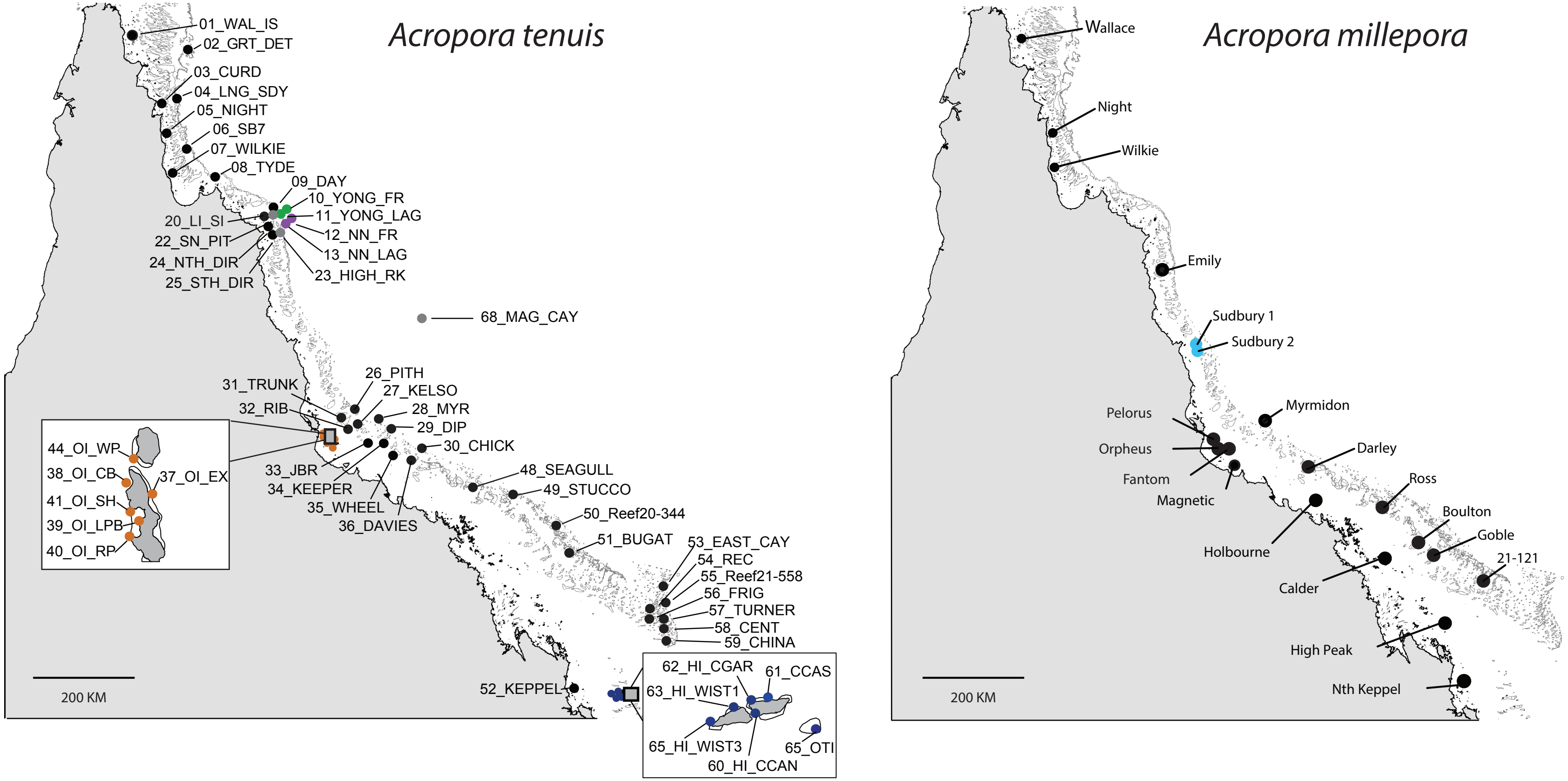


**Supplementary Figure 2:** Sampling locations for *Acropora tenuis* and *Acropora millepora*. Figures are modified from Lukoschek et al., 2016. Grey points indicate locations that were omitted for the present study and locations that were merged are shown in matching colours.

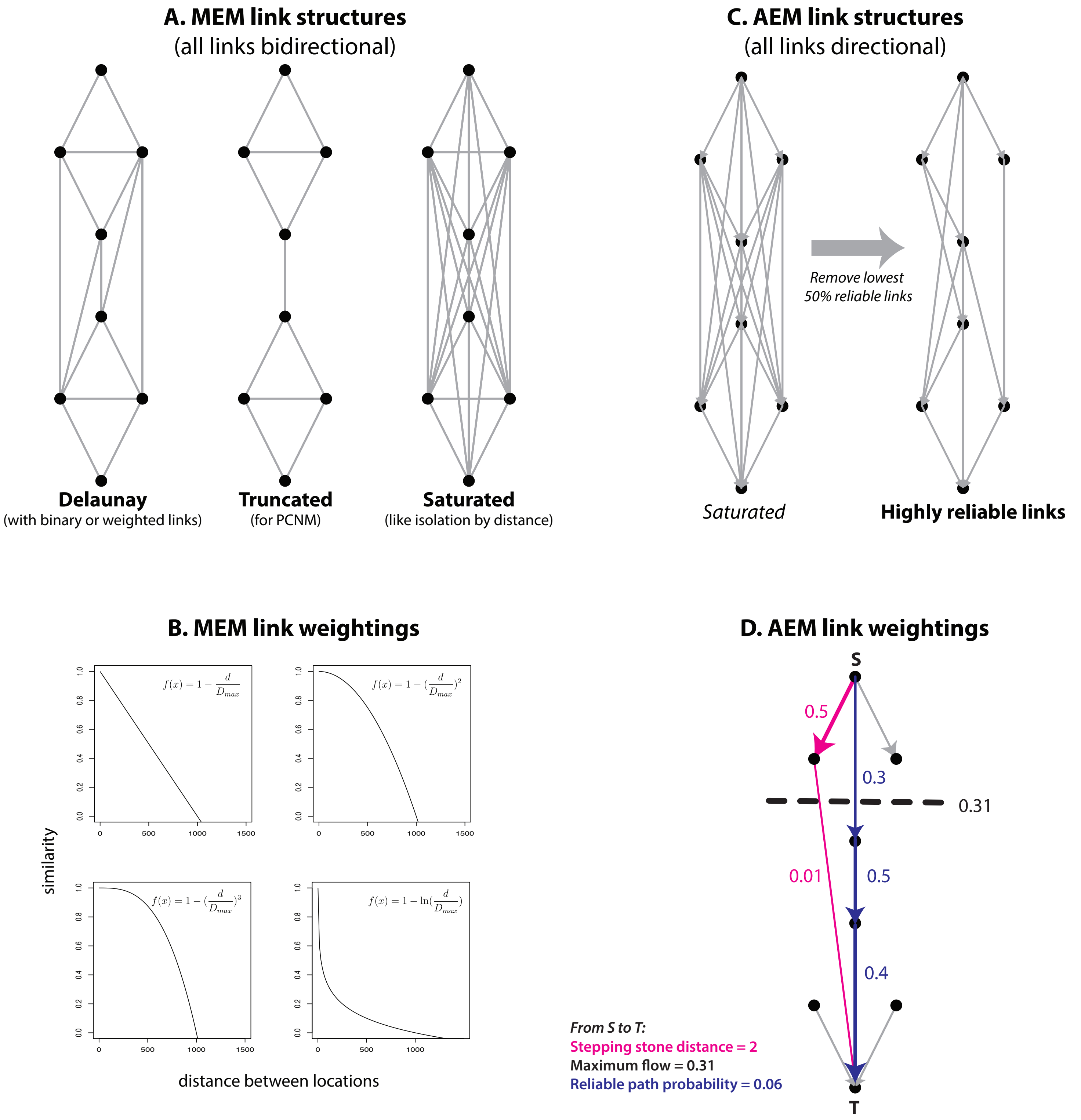


**Supplementary Figure 1**: Link structures and weighing schemes for spatial eigenvector models shown for an idealized set of sampling locations (black circles). Links are shown as grey lines. A) Moran Eigenvector Maps (MEMs) link structures depend solely on geographic position, where Delaunay triangulation and truncated criteria are commonly used to simplify the number of connections; connections between locations are bidirectional. B) MEM link weightings representing the strength of connections between locations can be unweighted (binary) or weighted where the increased influence of proximate locations is typically scaled by power functions (raised to the power of 1-3), and we also include a negative log scaling. C) Asymmetric Eigenvector Maps (AEMs) links are directional and indicated by arrowheads representing a directional process from top to bottom of the figure. Because the guidelines for constructing link structures for AEMs when sampling locations are irregular are not well established, we opted to simplify a saturated connection structure by only considering the more probable links using the reliable path criterion. D) AEM link weightings were based on three metrics: stepping-stone distance, maximum flow, and most reliable path probability. Considering two hypothetical locations S and T, the stepping-stone distance from S to T is the connecting path with minimum number of links, in this example a path of two steps shown in pink. The maximum flow is the sum of flow capacities through the minimum set of weakest connecting links that, if removed, would disconnect S from T, and the most reliable path is the product of link probabilities for the highest probability multistep pathway (shown in blue).

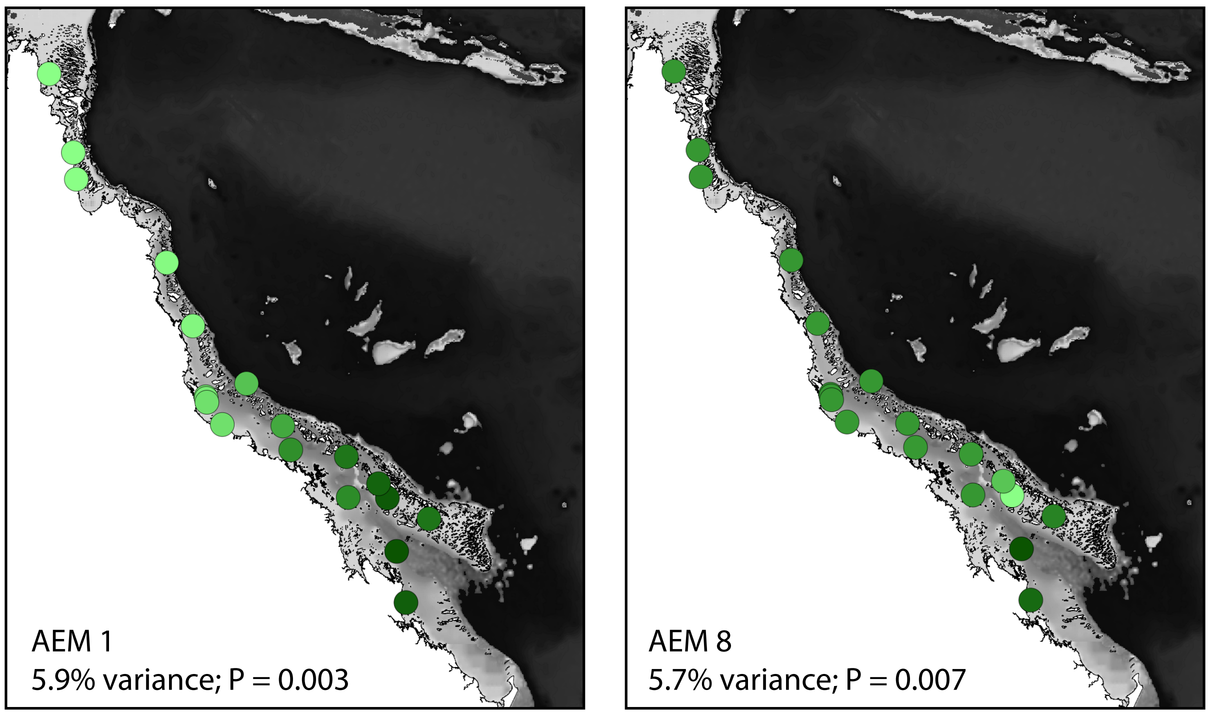


**Supplementary Figure 3:** Leading asymmetric eigenvector maps describing spatial genetic structure in *Acropora millepora*. Values by sampling location are colored by intensity of green hue. The greatest proportion of variance (AEM 1) describes a GBRF wide cline in genetic diversity followed by local scale spatial autocorrelation structures (AEM 8).

**Table S1:** Bounds for prior distributions of demographic model parameters. All parameters were drawn from a uniform distribution. A stepwise model of mutation was use to generate new alleles.

| Parameter | Lower Bound | Upper Bound |
|---|---|---|
| Effective population size ($Ne_i$) | 100 | 500000 |
| Migration rate ($m_{ij}$) | 0 | 0.5 |
| Time of divergence ($t_{divergence}$) | 5000 | 200000 |
| Mutation rate (μ) | $5 \times 10^{-7}$ | $5 \times 10^{-4}$ |

**Supplementary Table 2: Sampling locations for *Acropora tenuis***

| Original name | Reef ID in BP model | Lat Centroid | Long Centroid | Sample Size |
|---|---|---|---|---|
| 01_WAL_IS | 1 | -11.45 | 143.04 | 48 |
| 02_GRT_DET | 2 | -11.78 | 144 | 48 |
| 04_LNG_SDY | 3 | -12.51 | 143.79 | 29 |
| 03_CURD | 4 | -12.58 | 143.55 | 48 |
| 05_NIGHT | 5 | -13.18 | 143.57 | 48 |
| 31_TRUNK | 6 | -18.35 | 146.82 | 43 |
| 27_KELSO | 7 | -18.44 | 147 | 38 |
| 10_YONG_FR, 11_YONG_LAG | 8 | -14.61 | 145.63 | 51 |
| 12_NN_FR, 13_NN_LAG | 9 | -14.64 | 145.64 | 63 |
| 24_NTH_DIR | 10 | -14.74 | 145.51 | 46 |
| 08_TYDE | 11 | -13.98 | 144.52 | 45 |
| 25_STH_DIR | 12 | -14.82 | 145.53 | 48 |
| 06_SB7 | 13 | -13.4 | 143.97 | 48 |
| 09_DAY | 14 | -14.52 | 145.54 | 21 |
| 07_WILKIE | 15 | -13.77 | 143.64 | 47 |
| 20_LI_SI | 16 | -14.7 | 145.46 | 43 |
| 35_WHEEL | 17 | -18.8 | 147.53 | 44 |
| 36_DAVIES | 18 | -18.82 | 147.65 | 9 |
| 49_STUCCO | 19 | -19.56 | 149.6 | 19 |
| 32_RIB | 20 | -18.48 | 146.87 | 30 |
| 26_PITH | 21 | -18.21 | 147.02 | 44 |
| 28_MYR | 22 | -18.27 | 147.39 | 43 |
| 29_DIP | 23 | -18.41 | 147.45 | 19 |
| 37_OI_EX, 38_OI_CB, 39_OI_LPB, 40_OI_RP, 41_OI_SH, 41_OI_SH, 44_OI_WP | 25 | -18.59 | 146.49 | 268 |
| 48_SEAGULL | 27 | -19.53 | 148.98 | 21 |
| 33_JBR | 28 | -18.63 | 147.06 | 46 |
| 34_KEEPER | 29 | -18.75 | 147.72 | 49 |
| 30_CHICK | 30 | -18.66 | 147.71 | 20 |
| 50_Reef20344 | 31 | -20.78 | 150.9 | 45 |
| 51_BUGAT | 32 | -20.09 | 150.32 | 35 |
| 55_21558_LAG_F2 | 33 | -21.55 | 152.54 | 41 |
| 57_TURN_F1_F2_BK | 34 | -21.7 | 152.56 | 62 |
| 54_REC_F1_FR | 35 | -21.67 | 152.45 | 38 |
| 58_CENT_NTH_SW | 36 | -21.91 | 152.53 | 59 |
| 56_FRIG_FR_F1 | 37 | -21.73 | 152.44 | 41 |
| 53_EAST_SW_LAG | 38 | -21.5 | 152.56 | 49 |
| 60_HI_CCAN, 62_HI_CGAR, 62_HI_CGAR, 61_HI_CCAS, 61_HI_CCAS, 63_64_HI_WIS1, 65_HI_WIS3, 66_OTI_LAG, 67_OTI_WALL | 40 | -23.48 | 151.87 | 179 |
| 59_CHINA_SW_NTH | 42 | -22.01 | 152.65 | 51 |
| 52_KEPPEL | 43 | -23.17 | 150.93 | 13 |

### Supplementary Table 3: Sampling locations for *Acropora millepora*

| Original name | Reef ID in BP m | Lat Centroid | Long Centroid | Sample Size |
|---|---|---|---|---|
| Wallace | 1 | -11.4539 | 143.042 | 45 |
| Night_Is | 2 | -13.1837 | 143.5739 | 47 |
| Wilke | 3 | -15.606 | 145.6263 | 47 |
| Emily | 4 | -16.9984 | 146.205 | 96 |
| Sudbury_2, Sudbu | 5 | -13.7715 | 143.6402 | 47 |
| Myrmidon | 6 | -18.2657 | 147.3842 | 44 |
| Pelorus_SE | 7 | -18.5564 | 146.4889 | 50 |
| Orpheus_NE | 8 | -18.5933 | 146.4984 | 50 |
| N_Fantom | 9 | -19.1664 | 146.8504 | 49 |
| Magnetic_Is | 10 | -18.6767 | 146.5116 | 50 |
| Darley_Rf | 11 | -19.8727 | 149.5813 | 33 |
| Holbourne_Is | 12 | -19.725 | 148.3573 | 48 |
| Ross_Rf | 13 | -19.1969 | 148.1833 | 48 |
| Boulton_Rf | 14 | -20.7817 | 150.4846 | 51 |
| Calder_Is | 15 | -20.7717 | 149.6204 | 44 |
| Goble_Rf | 16 | -20.468 | 150.2905 | 51 |
| 21-121_Rf | 17 | -21.9571 | 150.6876 | 27 |
| High_Peak_Is | 18 | -21.2404 | 151.3989 | 44 |
| Nth_Keppel_Is | 19 | -23.0824 | 150.8929 | 47 |
